# Nonlinear modulation of human exploration by distinct sources of uncertainty

**DOI:** 10.1101/2025.04.16.649221

**Authors:** Xinyuan Yan, Becket R. Ebitz, David P. Darrow, Alexander B. Herman

## Abstract

Decision-making in uncertain environments requires balancing exploration and exploitation, with exploration typically assumed to increase monotonically with uncertainty. Challenging this prevailing assumption, we demonstrate a more complex relationship by decomposing environmental uncertainty into volatility (systematic change in reward contingencies, learnable) and stochasticity (random noise in observations, unlearnable). Across two behavioral experiments (N=1001, N=747) using a probabilistic reward task, we find a robust U-shaped relationship between the volatility-to-stochasticity (*v*/*s*) ratio and exploratory behavior, with participants exploring more when either stochasticity or volatility dominates. Remarkably, this pattern extends to real-world financial behavior, as demonstrated through analysis of five years of S&P 500 stock market data, where portfolio diversity (a proxy for exploration) shows the same U-shaped relationship with market volatility (systematic price movements driven by fundamental factors, e.g., economic shifts) relative to trading noise (random fluctuations from trading activity unrelated to fundamentals). These findings reveal how humans adaptively modulate exploration strategies based on the qualitative composition of uncertainty, with optimal performance occurring at intermediate uncertainty ratios. This nonlinear relationship has important implications for understanding decision-making across domains where uncertainty arises from multiple sources.

## Introduction

A fundamental tension lies at the heart of nearly every decision: persist with the current course or deviate to explore alternatives – the enduring exploration-exploitation dilemma (1). This dilemma exists because of environmental uncertainty, a pervasive feature of our surroundings (2). While substantial evidence indicates that uncertainty generally influences exploratory behavior (3, 4), less attention has been paid to how different types of uncertainty might have distinct effects.

Environmental uncertainty stems from multiple sources, with two critical dimensions being volatility and stochasticity (5). This distinction is crucial for understanding adaptive decision-making processes. For example, in medical diagnosis, distinguishing between volatility (changing disease prevalence due to seasonal patterns) and stochasticity (random variations in test results) can dramatically affect optimal treatment decisions. In reinforcement learning, these distinctions are also critical; highly volatile environments require higher learning rates to track changing values, while highly stochastic environments warrant lower learning rates to avoid overreacting to noise (6). Similarly, in financial markets, misattributing random price fluctuations (stochasticity) to fundamental changes in asset value (volatility) leads to excessive trading and diminished returns (7).

Intuition suggests opposing relationships between exploration and different uncertainty types. When volatility dominates, exploration should increase, as it helps track changing environments (5). Conversely, when stochasticity dominates, exploration theoretically offers minimal learning benefits since patterns are inherently unpredictable. However, real-world observations challenge this linear prediction. Even in highly stochastic environments, people actively diversify their choices rather than remaining static (8). This suggests a potentially U-shaped relationship: exploration should increase at both extremes of the volatility-stochasticity spectrum but for different functional reasons—as an environmental tracking mechanism under high volatility and as a risk-hedging strategy under high stochasticity. In financial markets, where success depends on finding signal in noise, under (9) and over (10) diversification are both associated with diminished performance. Notwithstanding the common wisdom about the importance of portfolio diversity, financial stability at the institutional level indeed appears to have a U-shaped relationship with diversification (11).

Despite the impact of different sources of uncertainty on behavior, separating them has been challenging. However, recent advances in computational modeling (5, 12) now provide quantitative tools to parse behavioral signals related to both volatility and stochasticity. Here, we utilize these time-series analysis tools to examine how exploratory behavior is shaped by the perceived structure of environmental uncertainty, operationalized as the volatility-to-stochasticity ratio. Through a combination of behavioral experiments and real-world financial market analysis, we discovered a U-shaped relationship: exploration increases both when volatility dominates and when stochasticity dominates.

## Results

### U-shaped relationship between *v*/*s* and exploration and its replication

We first examined the relationship between volatility and stochasticity (*v*/*s*) and exploration in two experiments (Experiment1, N=1001; Experiment2, N=747) employing a three-armed restless bandit task (13). Participants chose between three options on each trial, receiving probabilistic rewards based on underlying reward probabilities that changed independently and stochastically throughout the task (Method S1). We fitted participants’ choice data with a Kalman filter model (Method S2) to estimate their perceived volatility and stochasticity about the environment. Exploration was quantified as the rate (p(switch)) of choosing an option different from the last trial, a measure widely used to operationalize exploratory behavior in sequential decision-making tasks (14, 15)

The results supported the counterintuitive hypothesis, we found a significant U-shaped relationship between *v*/*s* and exploration (Figure 1A). Polynomial regression confirmed a significant positive quadratic term, indicating a non-linear relationship (Exp 1: b_quadratic_ = 0.005, t(998) =5.966, p < 0.0001). Intriguingly, there was an inverted-U shape between *v*/*s* and performance (Figure 1B) (Exp 1: b_quadratic_ = −0.002, t(998) = −6.938, p < 0.0001). We replicated our findings in another independent experiment (Figure 1C-D, Experiment 2, Table S1) with a fixed hazard rate of 1 and a step size of 0.1 for reward probability changes. U shape (Exp 2: *v*/*s* and p(switch): b_quadratic_ = 0.008, t(744) =10.033, p < 0.0001; *v*/*s* and performance: b_quadratic_ = −0.002, t(744) = −8.245, p < 0.0001).

**Figure 1.**
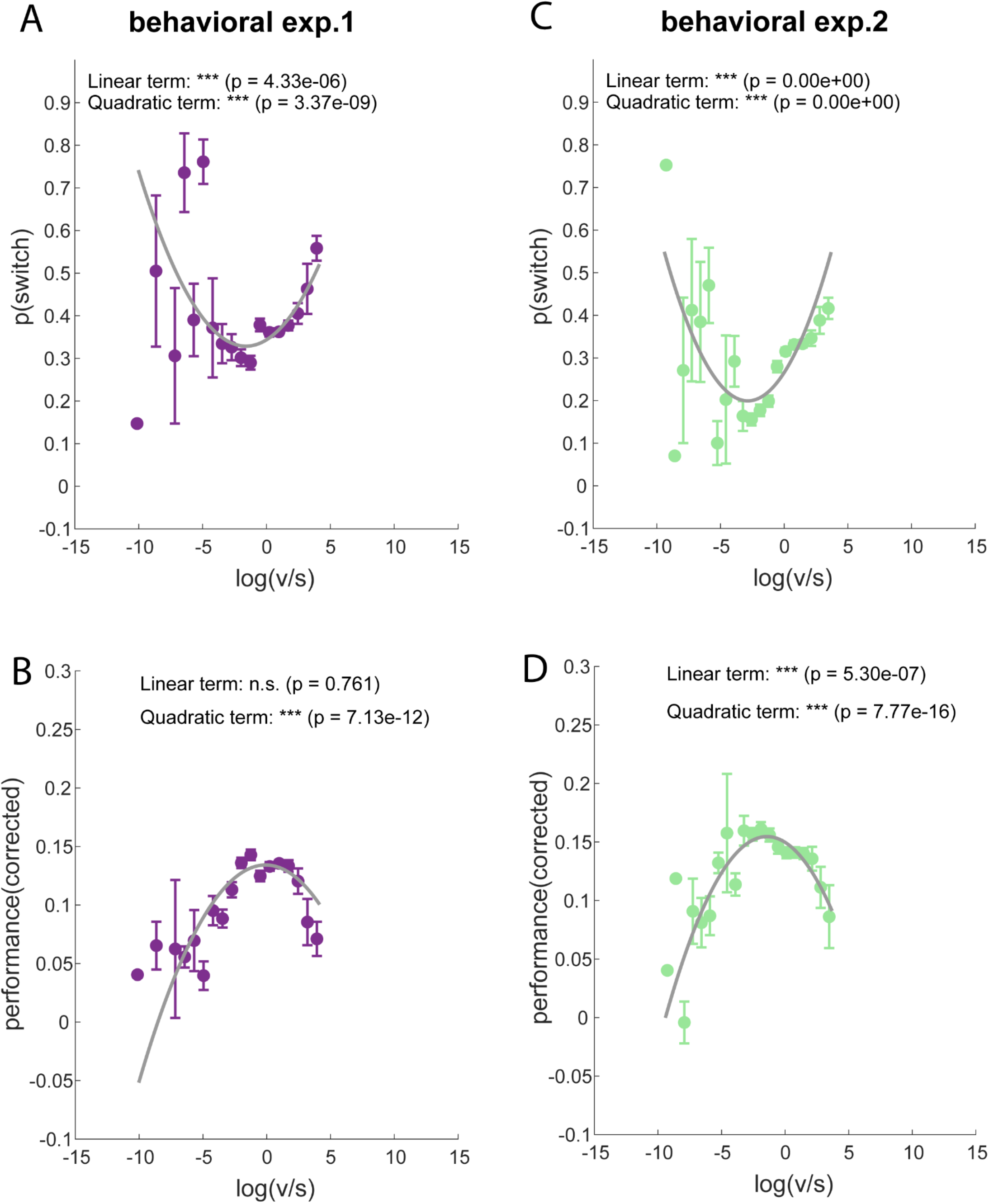
U-shaped relationship between volatility-stochasticity ratio and switching rate, performance in behavioral experiments. (A) U-shaped relationship between volatility-stochasticity ratio and exploration probability (p(switch)) in behavioral experiment 1 (N = 1001) (B) inverted-U shaped relationship between volatility-stochasticity ratio and performance (corrected by subtracting rewards at chance level) in behavioral experiment 1 (C) Replication of the U-shaped relationship between the volatility-stochasticity ratio and exploration in behavioral experiment 2 (N = 747) with different uncertainty settings. (D) Replication of the inverted-U shaped relationship between volatility-stochasticity ratio and performance in behavioral experiment 2. Data points (colored circles) represent binned averages for y-axis, where data were binned into 20 equal-width intervals along the x-axis, with error bars indicating standard error of the mean (SEM). The gray curve shows the quadratic regression fit. ***p < 0.001, n.s., non-significant.

The corresponding inverted U-shaped relationship between *v*/*s* and performance indicates that intermediate levels of perceived volatility relative to stochasticity may create optimal conditions for decision-making.

### Real-world data analysis

To assess the generalizability of these findings, we analyzed five years of daily S&P 500 data, focusing on the top 50 most actively traded stocks (see Method S7). We calculated the *v*/*s* ratio using a rolling window approach (window sizes: 100-150 days, step: 10 days), with volatility estimated as the rolling standard deviation of daily returns (16, 17) and stochasticity estimated using a composite measure of dispersion, autocorrelation, and a runs test (18) (see Method S2). We used portfolio diversity (1-Herfindahl-Hirschman Index of portfolio weights) (19, 20) as a proxy for exploration and average daily return as a measure of performance.

Remarkably, we observed the same U-shaped relationship between *v*/*s* and exploration in the S&P 500 data (Figure 2A). Polynomial regression confirmed a significant positive quadratic term (b_quadratic_ = 0.027, t(1084) =14.707, p < 0.0001). Furthermore, we found an inverted U-shaped relationship between *v*/*s* and performance (Figure 2B), with a significant negative quadratic term (b_quadratic_ = −0.003, t(1084) = −9.062, p < 0.0001). These findings demonstrate that the non-linear modulation of exploration and performance by *v*/*s* extends to real-world market behavior.

**Figure 2.**
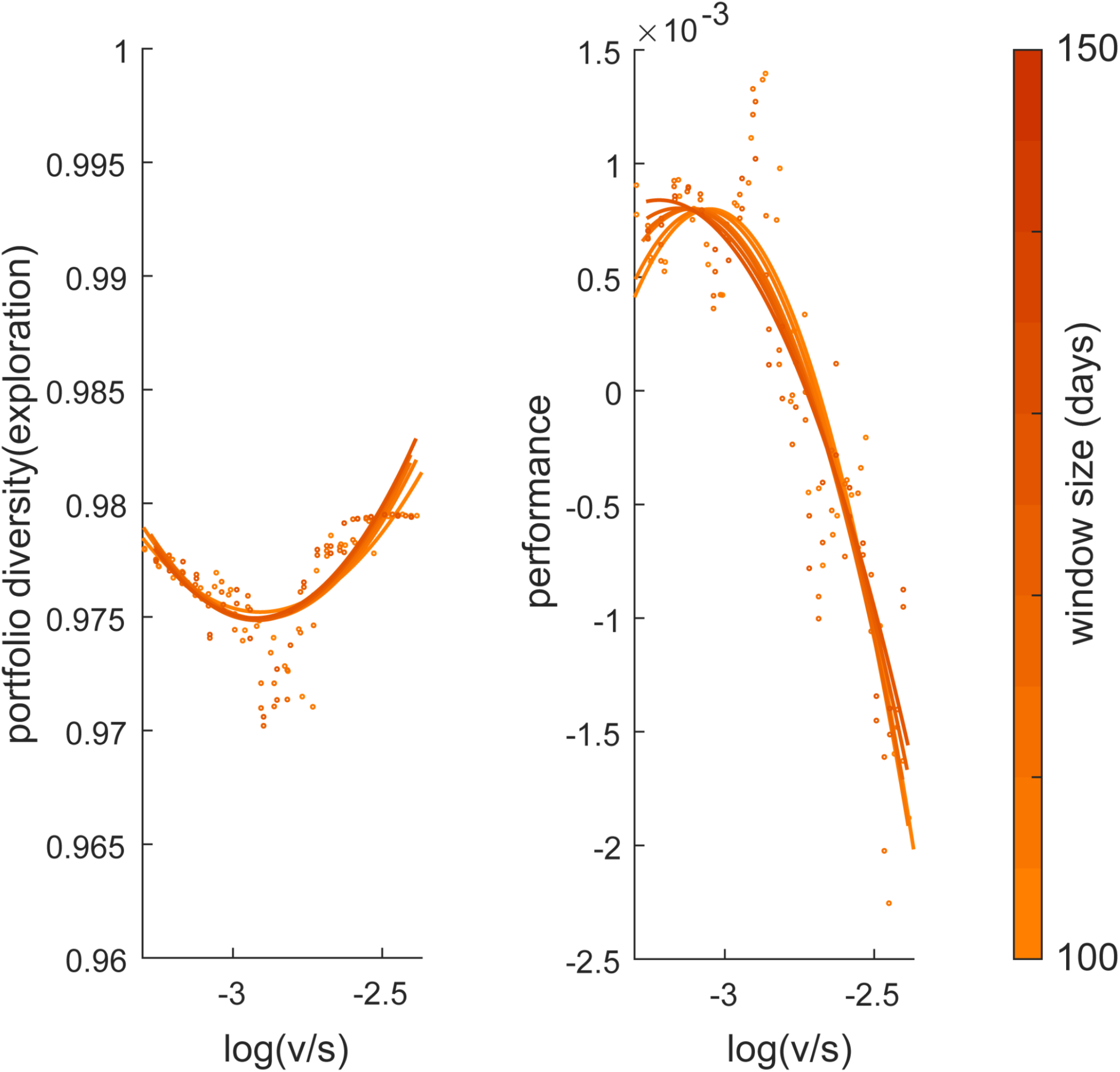
U-shaped relationship between volatility-stochasticity ratio and exploration, performance in S&P 500 market data. (A) the U-shaped relationship between exploration (portfolio diversity) and vol/stc for the S&P 500 data, and (B) the inverted U-shaped relationship between performance (average daily return) and vol/stc. Both relationships demonstrate that market behavior exhibits optimal exploration and performance at intermediate levels of the v/s ratio, consistent with our theoretical predictions. Data points (circles) represent binned averages for the y-axis, where data were binned into 20 equal-width intervals along the x-axis. The colored curves show the quadratic regression fits for each window size, with the color spectrum (light to dark orange) representing increasing window sizes (separate plots for each window size see Figure S1)

## Discussion

### Possible mechanisms driving exploration at different v/s ratios

Why does exploration increase at both ends of the v/s range? At low v/s ratios where stochasticity dominates, exploration may function as a risk-spreading strategy, evidenced by the higher switching rates even after rewards (Figure S2B and S2D). When observations are highly noisy, the expected value of any single option becomes difficult to estimate reliably. In such environments, distributing choices across multiple options could mitigate risk by avoiding overcommitment to options that appear favorable due to random fluctuations. At high v/s ratios, where volatility dominates, exploration likely serves as an environmental tracking mechanism (21), consistent with the increased switching after non-rewards (Figure S2A and S2C). Rapidly changing reward contingencies would incentivize continuous sampling of different options to detect and adapt to shifts in underlying probabilities. Future research should systematically examine these possible mechanisms, as well as their underlying neural correlates.

### Sweet spot: optimality at intermediate v/s ratios

The inverted U-shaped relationship between the v/s ratio and performance indicates that intermediate uncertainty ratios create optimal conditions for decision-making. By balancing these factors, individuals can effectively distinguish meaningful patterns from random noise without being overwhelmed by excessive environmental change, appropriately calibrating their exploration-exploitation trade-off.

### Consistent findings across internal and external uncertainty perception

Our findings indicate that a similar U-shape characterizes both the perceived uncertainty ratio of individuals (Exp.1) and the objective uncertainty ratio measured from external environments (Exp.2 and market data). This consistency suggests that the nonlinear relationship between exploratory behavior and the structure of uncertainty is a fundamental phenomenon, reflected not only in individual decision-making but also in aggregated macro-level market behavior.

### Future directions

Our findings contribute to understanding how different forms of uncertainty influence human decision-making and exploratory behavior across various contexts, including financial markets, psychology, computational psychiatry and reinforcement learning.

## Methods and Materials

### Exp.1

We reused our published dataset from (13). 1001 participants completed the three-armed restless bandit task (age range 18-54, mean ± SD = 28.446 ± 10.354 years; gender, 493 female). Task descriptions and detailed computational models are provided in Method S1.

### Exp.2

We recruited participants (non-clinical sample) via Prolific (Prolific. co); exclusion criteria included current or history of neurological and psychiatric disorders. 747 participants completed all questionnaires and the bandit task (age range 18-77, mean ± SD = 38.625 ± 12.757 years; gender, 370 female; Table S1). Task descriptions and detailed computational models are provided in Method S1-Method S6.

### Real-world market analysis (S&P 500)

We analyzed daily trading data from the S&P 500 stock market over five years to test our hypotheses in a real-world financial context. From the “all_stocks_5yr.csv” dataset (https://www.kaggle.com/datasets/rohitjain454/all-stocks-5yr), we first identified the 50 most actively traded stocks based on mean trading volume. Detailed experimental methods are provided in Method S7.

## Acknowledgments

This work was supported by the National Institute of Mental Health (Grant No. R21MH127607), the National Institute on Drug Abuse (Grant No. K23DA050909, to ABH), the University of Minnesota’s MnDRIVE (Minnesota’s Discovery, Research and Innovation Economy) initiative (to DPD and XY). We sincerely thank Joshua Blumenstock for helpful comments on the manuscript.

## SUPPLEMENTARY INFORMATION

### Method S1. Three-armed restless bandit task in Exp.1 and Exp.2

#### Exp.1

Participants (N=1001) were free to choose between three targets for the potential to earn a reward of 1 point. Each target is associated with a hidden reward probability that randomly and independently changes throughout the task. We seeded each participant’s reward probability randomly to prevent biases due to particular kinds of environments. Specifically, on each correct trial, there was a 67% chance that the reward probability for each target would either increase or decrease by 0.2, with these probabilities bounded between 0 and 1.0. Due to the variable and independent nature of the rewards, participants could only estimate the probabilities by actively sampling from the targets and accumulating their reward experiences over time.

#### Exp.2

The Procedure was the same as in Exp.1, but on each correct trial, there was a 100% chance that the reward probability for each target would either increase or decrease by 0.1, with these probabilities bounded between 0 and 1.0.

### Method S2. Kalman filter model

The Kalman filter (KF) model has been widely applied in psychology and neuroscience to study various aspects of learning and decision-making (1, 2). Details to explain the kalman filter model are provided below.

In the Kalman filter model for a multi-armed bandit task, *process noise* and *observation noise* refer to two distinct sources of uncertainty that affect the learning and decision-making process.

Process noise represents the uncertainty in the evolution of the hidden state (reward mean) over time. It accounts for how the true state evolves from one point in time to the next. In mathematical terms, process noise is part of the state transition equation in the Kalman Filter:

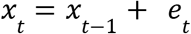

*x* _*t*_ is the state at time *t*

*e*_*t*_ is the process noise *t*, which is assumed to be drawn from a normal distribution with zeros mean and process noise variance *v*. Where the *e*_*t*_∼*N*(0, *v*).

The process noise captures the idea that the reward-means for each arm can change from one trial to the next, even in the absence of any observations. A higher process noise variance *v* indicates a more volatile environment, where the reward means are expected to change more rapidly.

In contrast, observation noise represents the uncertainty in the observed rewards, given the current hidden state (reward mean). Which is assumed to be Gaussian with zero mean and a fixed variance σ^2^.

The observation noise captures the idea that the observed rewards are noisy and can deviate from the true reward mean due to random fluctuations or measurement errors. A higher measurement noise variance indicates a more stochastic environment, where the observed rewards are less reliable and informative about the underlying reward means.

The Kalman Filter operates optimally when the statistical properties of the process noise and the measurement noise are accurately known.

When observation noise variance (σ^2^) is high relative to the process noise variance (*v*), the Kalman gain will be small, and the model will rely more on its prior beliefs and less on noisy observations. Conversely, when the observation noise variance (*v*), is high relative to the process noise variance (σ^2^), the Kalman gain will be large, and the model will update its beliefs more strongly based on the observed rewards.

#### Extended Kalman filter for three-armed bandit task

The Kalman filter model can be extended to capture the effects of both volatility and stochasticity in a multi-armed bandit task (3, 4).

In the current study, process noise variance (*v*) and observation noise variance (σ^2^) represent volatility and stochasticity, respectively.

A traditional assumption of the Kalman filter is that the process noise variance, *v*, as well as the observation noise variance, σ^2^are constant.

Reward means update:

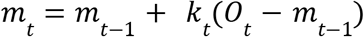

Where *m* _*t*_ is the estimated mean or value of the chosen arm at time *t* and *O* _*t*_ is the observed reward at time *t*.

The mean update is driven by the prediction error, which is the difference between the observed reward and the previous estimate.

Kalman gain is defined as:

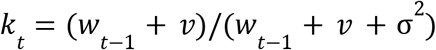

Here, *k* _*t*_ represents the Kalman gain or learning rate, which adjusts the weight given to new information based on the relative uncertainty of the prior estimate (*w* _*t*−1_) and the total noise (*v* + σ^2^). When the stochasticity (σ^2^) is high relative to the volatility (*v*), the Kalman gain (learning rate) will be small, and the model will rely more on its prior beliefs and less on the observations. Conversely, when the volatility (*v*), is high relative to the stochasticity (_σ_^2^), the Kalman gain (learning rate) will be large, and the model will update its beliefs more strongly based on the observed rewards.

Variance update equation:

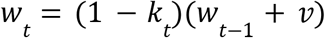

This equation updates the posterior variance (*w* _*t*_), which represents the estimate’s uncertainty after observing *O* _*t*_.

### Method S3. Alternative model

We also fitted our data with alternative models including a volatile kalman filter model

#### Volatile kalman filter for three-armed bandit task

The key difference between a standard Kalman filter and a volatile Kalman filter (VKF) is the variance of the process noise, a stochastic variable that changes with time. In other words, the VKF introduces parameters to handle the volatility in the process noise. Specifically, it allows the process noise variance *v* to vary with the observed prediction errors, reflecting changes in environmental volatility.

Our approach here is essentially the same as that taken by Piray and Daw (4). Here, we briefly described the model details as follows.

Kalman gain:

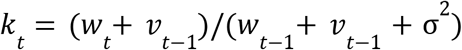

Update for the reward means:

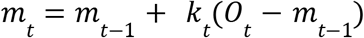

Update for posterior variance *w* _*t*_:

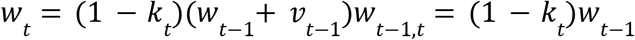

Update for volatility:

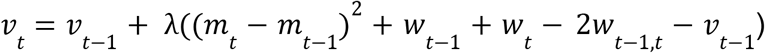

### Method S4. Model comparison

For model comparison, we used Bayesian information criterion (BIC) to select winning models, BIC values can be found in Table S2.

### Method S5. Model validation

We validated our modeling procedure using two approaches. First, we tested the accuracy of the model prediction. We calculated the correlation between behavioral output predicted by the model and real choices. We conducted this analysis with a randomly selected subset of 50 participants from the full dataset of 1001 and demonstrated strong correlations between observed behaviors and model predictions across three cues: r = 0.719 for cue1, r = 0.730 for cue2, and r = 0.775 for cue3. All correlations were statistically significant, p < 0.0001; see Figure S3)

Secondly, we conducted parameter recovery analyses using synthetic data generated from the Kalman Filter model (Method S2). For each simulation, we generated data for 100 agents, with each subject completing three sequences of 300 trials (3 different cues). We obtained reasonable parameter recovery correlations. The mean Pearson correlations were 0.707, 0.671, 0.973 for *v*, σ2, and β, respectively (Figure S4)

### Method S6. Split-half reliability

To assess the split-half reliability of our task, we examined the consistency of overall choices and model parameters from the winning model between the first and second halves of trials. For overall choice proportion, we employed Pearson’s correlations to calculate reliability. For model parameters, we utilized a more sophisticated approach, calculating model-derived estimates of Pearson’s r values from the parameter covariance matrix. This method, which estimates first- and second-half parameters within a single model, has recently been validated for accurate parameter reliability estimation (5). We interpreted indices of reliability based on conventional values of <0.40 as poor, 0.4–0.6 as fair, 0.6–0.75 as good, and >0.75 as excellent reliability (Fleiss, 2011). Overall choice proportion showed fair-to-good reliability (r=0.65, r=0.64, r=0.53 for *v*, σ^2^, and β, respectively). The model parameters showed good-to-excellent reliability (r = [0.79, 0.78, 0.69] after Spearman-Brown correction) (Figure S5).

### Method S7. Real-world market analysis (S&P 500)

#### Data selection and preprocessing

We analyzed daily trading data from the S&P 500 stock market over five years to test our hypotheses in a real-world financial context. From the “all_stocks_5yr.csv” dataset (https://www.kaggle.com/datasets/rohitjain454/all-stocks-5yr), we first identified the 50 most actively traded stocks based on mean trading volume. This selection process focused our analysis on the most liquid securities while maintaining a diverse cross-section of the market.

All timestamps were converted to a standardized datetime format. We constructed a price matrix (time × stocks) containing daily closing prices for each selected security across all trading days in the sample period. To ensure data integrity, we systematically identified missing values (0.05% of observations) and imputed them using linear interpolation, preserving the temporal structure of the price series.

#### Returns calculation and time series construction

Daily returns were computed for each stock *i* as:

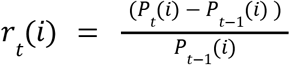

where *P* _*t*_ (*i*) represents the closing price. This transformation yielded a returns matrix R ∈ ℝ ^*T*×50^, where T denotes the number of trading days and N = 50 represents the number of stocks.

#### Rolling window analysis

To capture the temporal dynamics of market uncertainty, we implemented a rolling window approach with systematic variation in window size. Specifically, we analyzed window sizes ranging from 100 to 150 trading days (approximately 5-7 months) in increments of 10 days. For each window size w and time point t, we calculated uncertainty metrics using return data from the interval [t-w+1, t].

#### Volatility estimation

For each rolling window, we quantified volatility as the moving standard deviation of returns:

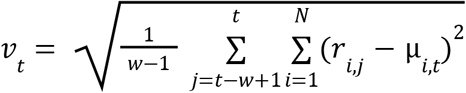

where μ _*i*,*t*_ is the mean return of stock i in the window ending at time t. This measure captures the overall dispersion of returns across all assets, providing a market-level assessment of process noise (volatility).

#### Stochasticity estimation

We developed a composite stochasticity metric that integrates three complementary dimensions of observational noise:

1. Return Dispersion (*D* _*t*_), which was defined by the standard deviation of all returns within the window, capturing the overall variability:

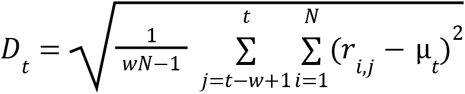
2. Temporal independence (*A* _*t*_), using 1 minus the mean absolute value of autocorrelations up to lag 5.

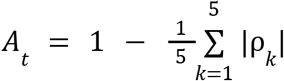
3. Runs test statistic (*R* _*t*_), which was defined by a normalized measure of the randomness in the sequence of returns.

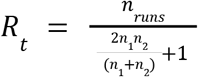

where *n*_*runs*_ is the number of runs in the binary sequence created by comparing returns to their mean value, and n_1_ and n_2_ are the counts of above-mean and below-mean returns, respectively. For example, if we have a sequence of returns and mark each value as “+” if it’s above the mean and “-” if it’s below the mean, we might get an sequence like: ++---+++--+

In this example, there are 5 runs: two consecutive “+”, three consecutive “-”, three consecutive “+”, two consecutive “-”, and one “+”. Values closer to 1 indicate greater randomness.

These three components were equally weighted to form the composite stochasticity measure:

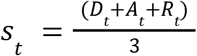

To ensure numerical stability, we applied Winsorization at the 1st and 99th percentiles and established a minimum threshold of machine epsilon (≈2.22 × 10^−16^) for all stochasticity values.

#### Volatility-stochasticity ratio

We calculated the log-transformed ratio of volatility to stochasticity: log(*v*/*s*). Data points with extremely small stochasticity values (< 2.22e-15) were filtered out to avoid numerical instability. Additionally, we removed outliers beyond the 1st and 99th percentiles of the ratio distribution, as well as extreme values where the ratio exceeded 100.

#### Exploration and performance metrics

To operationalize exploration in the financial context, we developed a portfolio diversity measure based on the Herfindahl-Hirschman Index (HHI). For each time window ending at t, we:

1. Calculated mean returns for each stock i within the window

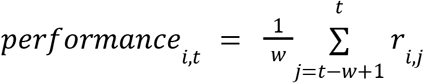
2. Transformed these performance metrics into non-negative weights by normalizing relative to the minimum performance:

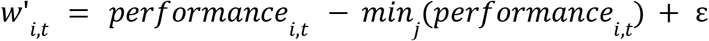

where ε is a small constant (≈2.22 × 10^−16^) to ensure strictly positive weights.
3. Standardized the weights to sum to unity

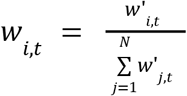
4. Computed portfolio diversity as the complement of the Herfindahl-Hirschman concentration index

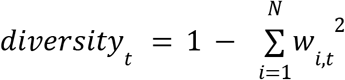

This measure ranges from 0 (complete concentration in a single asset, representing pure exploitation) to approximately 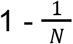 (perfect diversification across all assets, representing maximum exploration).

Higher diversity values indicate more distributed portfolio allocations, analogous to increased exploration in bandit decision-making contexts.

#### Performance assessment

We measured market performance using the average return across all stocks within each ***w***-day window.

#### Statistical analyses

We conducted polynomial regression analyses to examine the relationship between log-transformed *v*/*s* and both exploration and performance measures for each window:

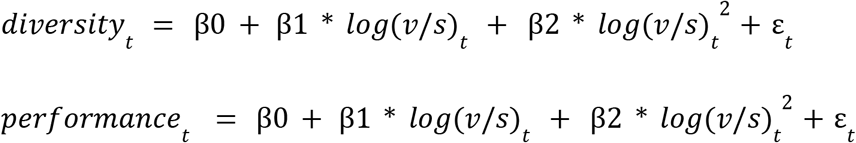

For visualization purposes, we binned the log-transformed ratios into equal intervals and computed mean portfolio diversity and performance metrics within each bin. The resulting relationships were plotted with quadratic fit overlays to illustrate the U-shaped and inverted U-shaped patterns for exploration and performance, respectively.

**TableS1.**
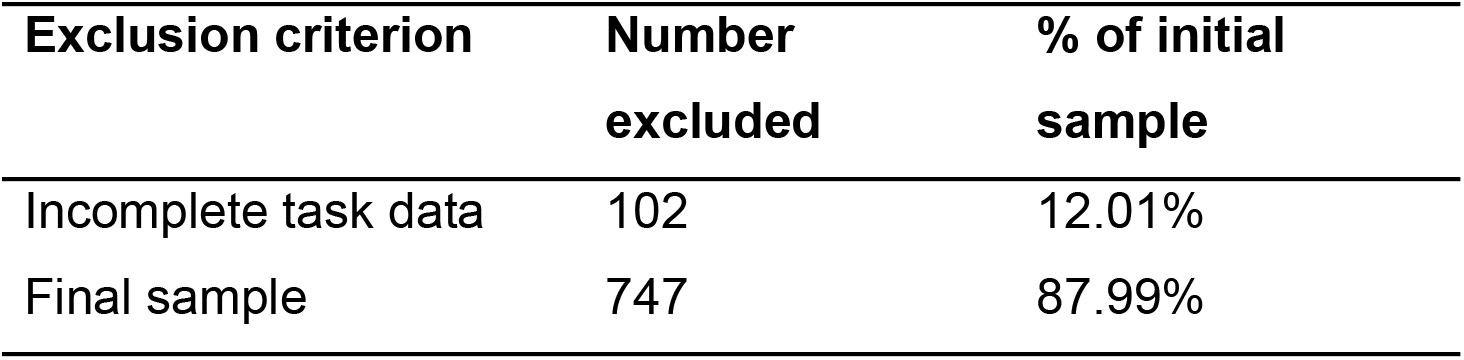
Detailed exclusion criteria table for Exp.2.

**Table S2.**
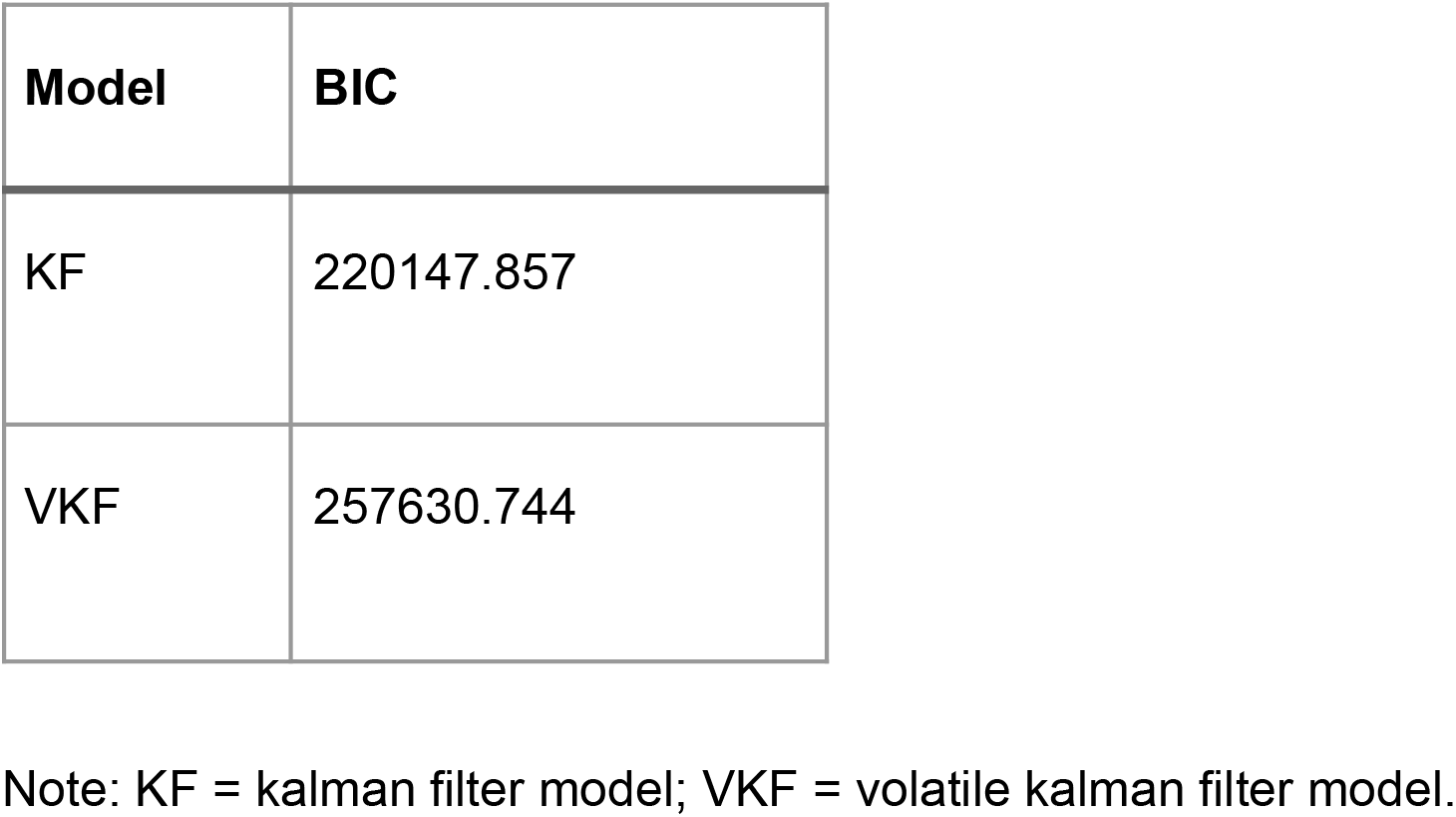
Model comparison.

| **Model** | **BIC** |
| --- | --- |
| KF | 220147.857 |
| VKF | 257630.744 |

Note: KF = kalman filter model; VKF = volatile kalman filter model.

**Figure S1.**
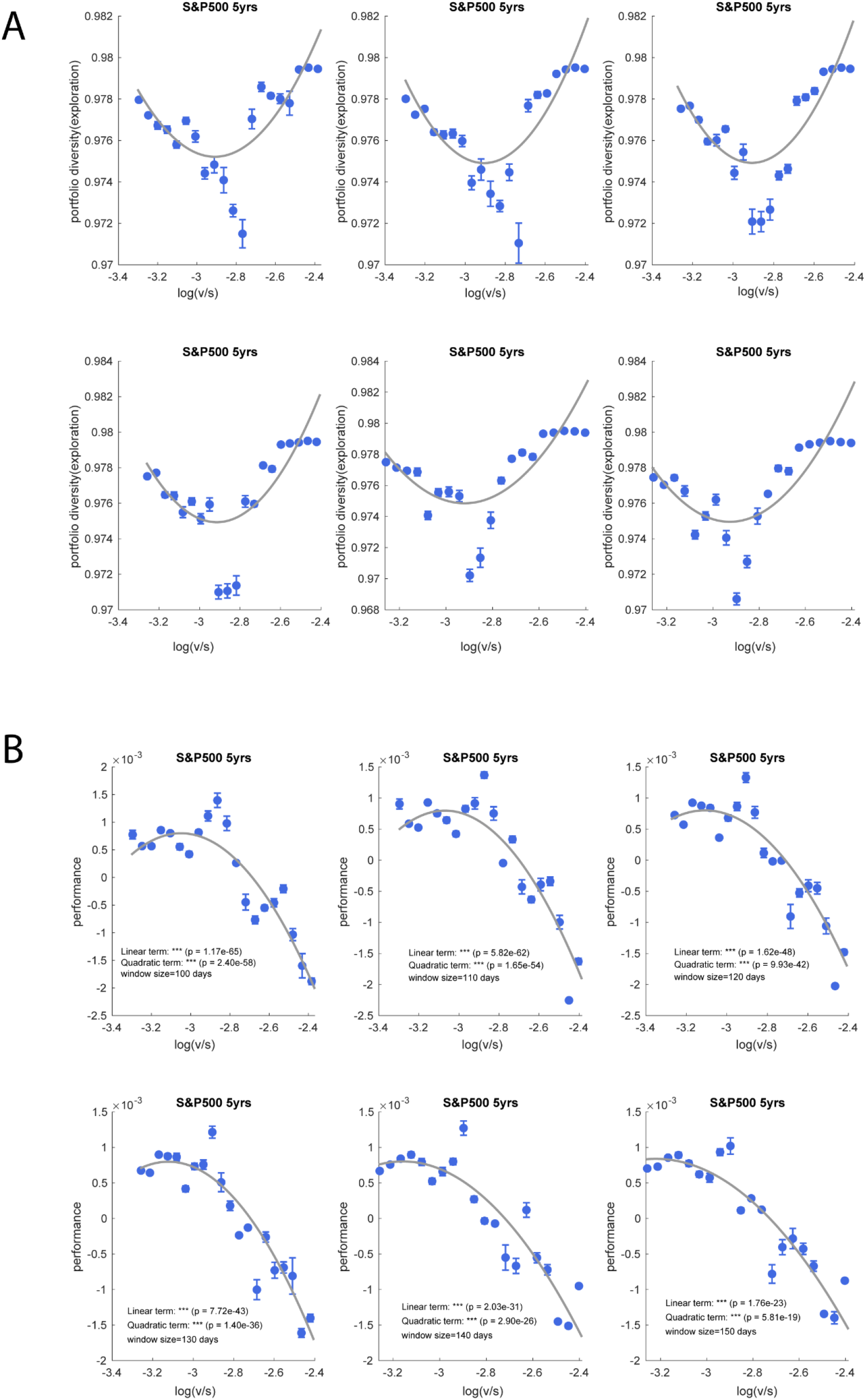
Non-linear relationship between volatility/stochasticity ratio and exploration/performance in S&P 500 market data. (A) U-shaped relationship between the volatility/stochasticity (v/s) ratio and portfolio diversity (exploration) across different rolling window sizes (100-150 days). Data points (circles) represent binned averages for portfolio diversity and performance, where data were binned into 20 equal-width intervals along the x-axis, with error bars indicating the standard error of the mean (SEM). Statistical annotations indicate the significance of the quadratic terms, with the significant positive quadratic terms (p < 0.001) confirming the U-shaped relationship. (B) Inverted U-shaped relationship between the v/s ratio and performance (average daily returns) across the same window sizes. The significant negative quadratic terms (p < 0.001) confirm the inverted U-shaped relationship. Both relationships demonstrate that market behavior exhibits optimal exploration and performance at intermediate levels of the v/s ratio, consistent with our theoretical predictions.

**Figure S2.**
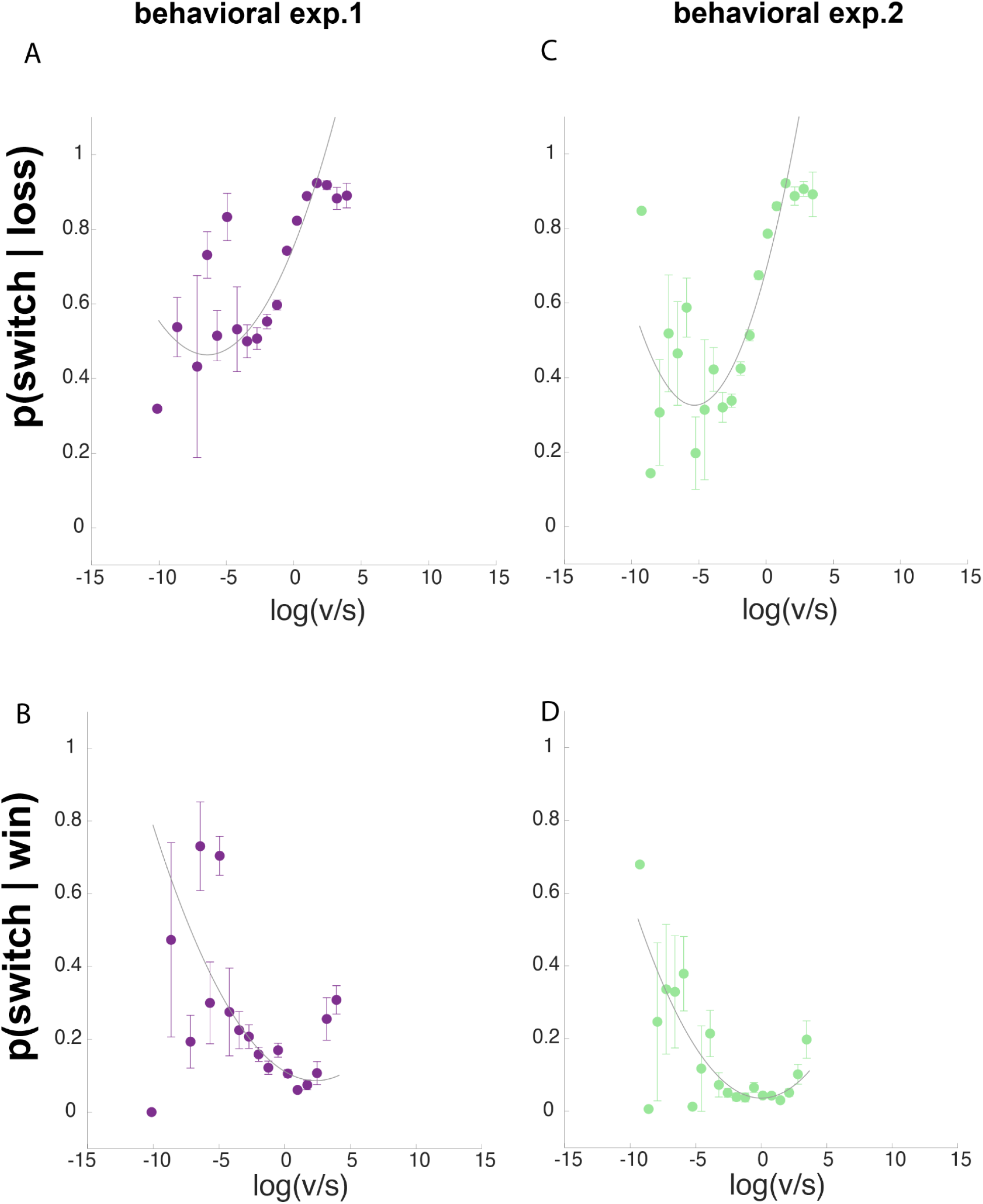
Human exploration strategies differ based on the volatility/stochasticity ratio A) In behavioral experiment 1, participants exhibited significantly higher switching rates following non-reward outcomes in high volatility/stochasticity (v/s) environments compared to low v/s environments. (B) Conversely, participants switched more frequently even after receiving rewards in low v/s environments than in high v/s environments. These patterns were successfully replicated in behavioral experiment 2. These results reveal that humans employ context-dependent exploration strategies: in high v/s conditions, exploration appears driven by loss avoidance behavior, while in low v/s conditions, exploration persists despite rewards, suggesting a risk-spreading strategy in response to perceived randomness in more stochastic, unstable environments.

**Figure S3.**
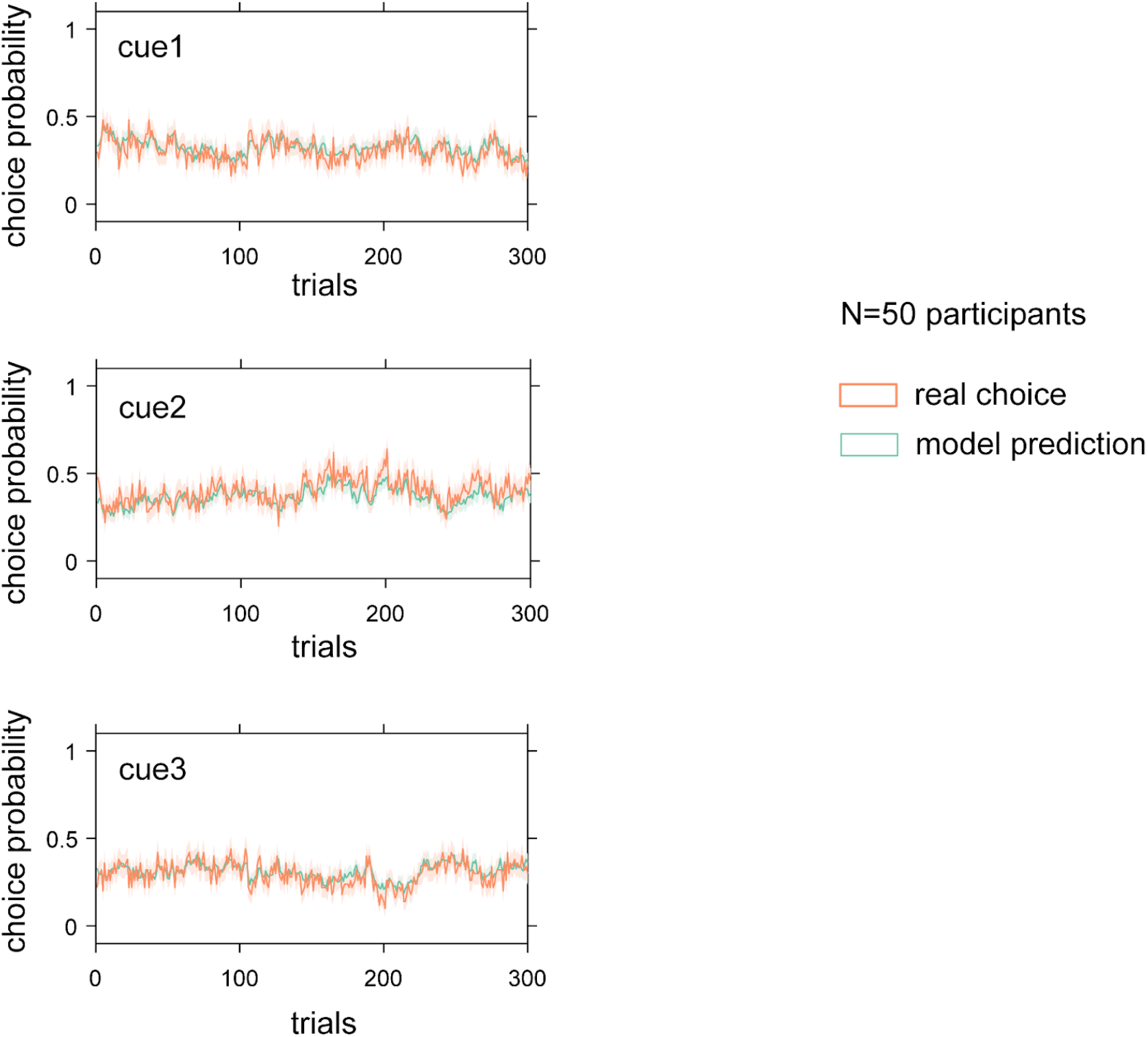
Posterior predictive check for model validity. This analysis was conducted with a randomly selected subset of 50 participants from the full dataset. It demonstrated strong correlations between observed behaviors and model predictions across three cues: r = 0.719 for cue1, r = 0.730 for cue2, and r = 0.775 for cue3. All correlations were statistically significant (p < 0.0001).

**Figure S4.**
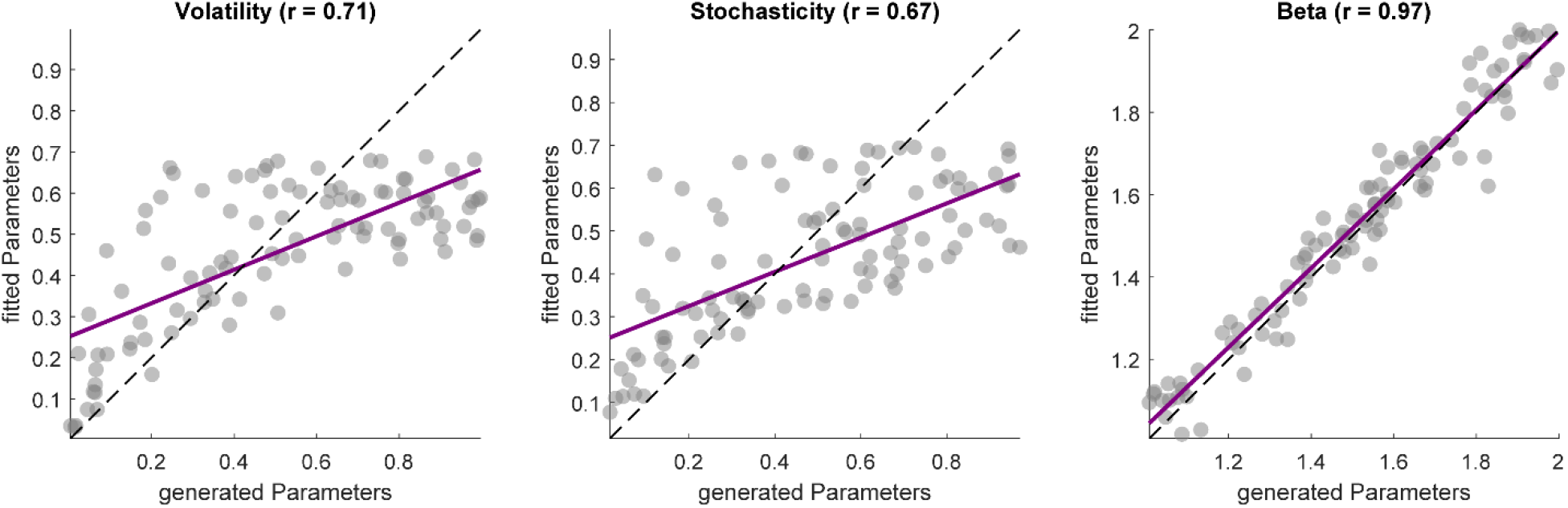
Parameter recovery. Following the standard procedure (4), we conducted parameter recovery analyses using synthetic data generated from the Kalman Filter model. For each simulation, we generated data for 100 agents, with each subject completing three sequences of 300 trials (3 different cues). We ran 50 simulations per agent and analyzed recovery using Pearson correlations between true parameters and averaged fitted parameters. We obtained reasonable parameter recovery correlations. Pearson correlations were for *v* = 0.707, σ^2^= 0.671, and β = 0.973.

**Figure S5.**
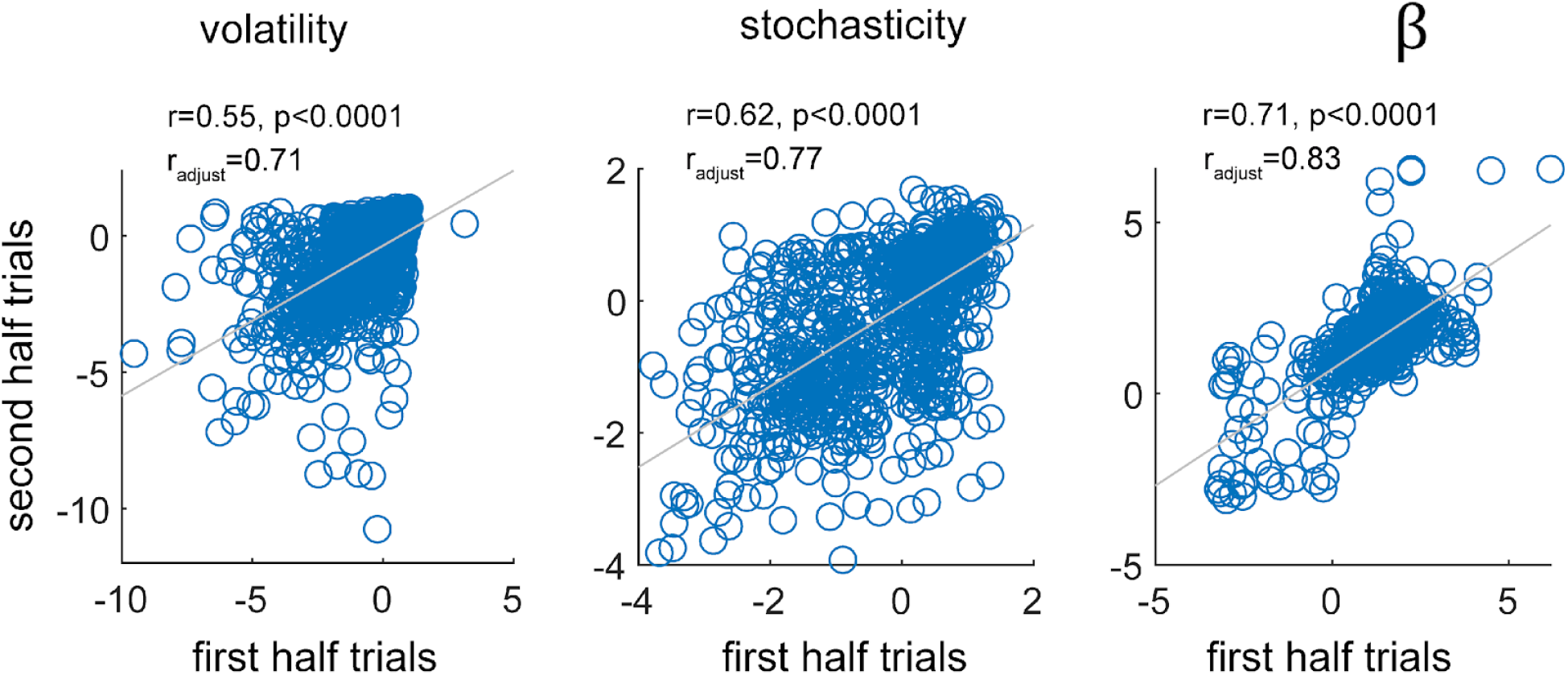
Split-half reliability of the task. We assessed the reliability of our task measures using a split-half approach. The scatter plots comparing parameters (before transformation) from the first and second halves of the task are presented, along with their corresponding reliability estimates (Pearson’s r values). Reliability estimates for the computational measures from the winning computational model were computed by fitting split-half parameters within a single model and then using the parameter covariance matrix to derive Pearson’s correlation coefficients for each parameter across halves. Reliability estimates are reported as unadjusted values (r) and after adjusting for reduced number of trials via Spearman-Brown correction (r_adjust_). Statistics reported here based on the correlation between transformed parameters between these two halves. Grey lines show lines-of-best-fit. Overall choice proportion showed fair-to-good reliability (r=0.55, r=0.62, r=0.71 for volatility, stochasticity, and β, respectively). The model parameters showed good-to-excellent reliability (r = [0.71, 0.77, 0.83] after Spearman-Brown correction).

